# The role of the γ subunit in the photosystem of the lowest-energy phototrophs

**DOI:** 10.1101/2022.09.24.509313

**Authors:** Dowrung Namoon, Nicola M Rudling, Daniel P Canniffe

**Affiliations:** Department of Biochemistry & Systems Biology, Institute of Systems, Molecular & Integrative Biology, University of Liverpool, UK

## Abstract

Purple phototrophic bacteria use a core ‘photosystem’ consisting of light harvesting antenna complex 1 (LH1) surrounding the reaction centre (RC), which primarily absorbs far-red–near-infrared light and converts it to chemical energy. Species in the *Blastochloris* genus, which are able to use light >1000nm for photosynthesis, use bacteriochlorophyll (BChl) *b* rather than the more common BChl *a* as their major photopigment, and also uniquely assemble LH1 with an additional polypeptide subunit, LH1γ, encoded by multiple open reading frames in their genomes. In order to assign a role to this subunit, we deleted the four LH1γ-encoding genes in the model *Blastochloris viridis*. Interestingly, growth under halogen bulbs routinely used for cultivation of anoxygenic phototrophs yielded cells displaying an absorption maximum of 825 nm, similar to that of the RC complex without LH1, but growth under white light from fluorescent bulbs yielded cells with an absorption maximum at 972 nm. HPLC analysis of pigment composition and sucrose density gradient fractionation demonstrate that the mutant grown in white light assembles RC–LH1, albeit with an absorption maximum blue-shifted by 46 nm relative to the WT complex. Wavelengths between 900–1000 nm transmit poorly through the atmosphere due to strong absorption by water, thus our results provide an evolutionary rationale for the incorporation of the γ subunit into the LH1 ring; this polypeptide red-shifts the absorption maximum of the complex to a range of the spectrum where the photons are of lower energy but are more abundant. Finally, we transformed the mutant with plasmids carrying genes encoding natural LH1γ variants and demonstrate that the polypeptide found in the WT complex red-shifts absorption back to 1018 nm, but incorporation of a distantly-related variant results in only a moderate red-shift. This result suggests that tuning the absorption maximum of this organism is possible, and may permit light capture past the current low-energy limit of natural photosynthesis.

## INTRODUCTION

Chlorophototrophic organisms use chlorophyll (Chl) and/or bacteriochlorophyll (BChl) pigments to capture solar radiation and convert it to chemical energy to power cellular metabolism [1]. Oxygenic chlorophototrophs, such as plants, algae and cyanobacteria, use Chls to primarily absorb photons of visible wavelengths, which have sufficient energy to drive the thermodynamically-challenging extraction of electrons from water, liberating molecular oxygen as a by-product [2]. Anoxygenic phototrophic bacteria are generally found below oxygenic organisms in water columns and microbial mats, and use BChls to capture lower-energy wavelengths in the far-red and near-infrared region of the spectrum that have not been utilised by the Chl-containing organisms above [3]. These bacteria use alternative sources of electrons than water, and thus do not generate O_2_. The vast majority of anoxygenic phototrophs use BChl *a* to harvest in the 780–900 nm range, while a small number of phototrophs found within the Proteobacteria use BChl *b*, and harvest wavelengths greater than 1000 nm [4].

BChl *b* was discovered in 1963 as the sole BChl extracted from an organism tentatively identified as a species of *Rhodopseudomonas*; the extracted pigment displayed a Q_y_ maximum red-shifted by 23 nm compared to that of BChl *a* [5]. An additional *‘Rhodopseudomonas’* isolate containing this pigment was found to have a whole-cell absorption maximum at ^~^1020 nm, 142 nm further into the near-infrared than the BChl *a*-containing proteobacterium *Rhodospirillum rubrum* to which it was compared [6]. This strain was subsequently named *Rhodopseudomonas viridis* due to its intense green colour [7]. Phylogenetic analysis of this strain, and closely-related species synthesising BChl *b*, led to transferral to the novel *Blastochloris* genus; the isolate described above being designated *Blastochloris (Blc.) viridis* [8].

The reaction centre (RC), the site of charge separation that initiates photosynthetic electron transfer, is encircled by an antenna known as light-harvesting complex 1 (LH1) to form the ‘core’ RC–LH1 supercomplex, the key functional unit for phototrophy in Proteobacteria. RC–LH1 complexes contain 5 universal components: the L, M, and H subunits of the RC, and the α and β polypeptides of LH1. Most proteobacterial RCs, including that of *Blc. viridis*, have a bound cytochrome subunit, C, containing 4 haem cofactors. The RC from *Blc. viridis* was the first membrane protein to have its structure solved, earning the 1988 Nobel Prize in Chemistry [9,10]. Some LH1 antennas also contain additional subunits; the *Rhodopseudomonas palustris* LH1 ring contains a single transmembrane helix known as protein W [11,12], LH1 from *Rhodobacter* spp. contains a PufX polypeptide [13–15], and a further subgroup of these organisms additionally contain PufY [16–18]. These polypeptides create a channel in the LH1 ring allowing quinone/quinol exchange between the RC and the cytochrome *bc*_1_ complex [19,20]. An additional LH1 subunit was also identified in *Blc. viridis* [21]. However, unlike those mentioned above, this LH1γ polypeptide was found to be in apparent equimolar ratio with the α and β polypeptides [22]. To date, no ortholog of the LH1 γ polypeptide is found outside of the *Blastochloris* genus. A recent cryo-electron microscopy structure of the RC–LH1 complex from *Blc. viridis* revealed the location of γ, packing between β polypeptides on the outside of the ring (**Fig. 1**) [16]. The α:β:γ ratio was found to be 17:17:16; in this case the ‘missing’ γ subunit creates the channel for quinone diffusion. We proposed that the role of the γ subunit in this complex is to tighten packing of the BChls in LH1, increasing excitonic coupling of these pigments, which results in the extreme ‘red-shift’ of the complex past 1000 nm to the current red limit for photosynthesis on Earth.

**Figure 1.**
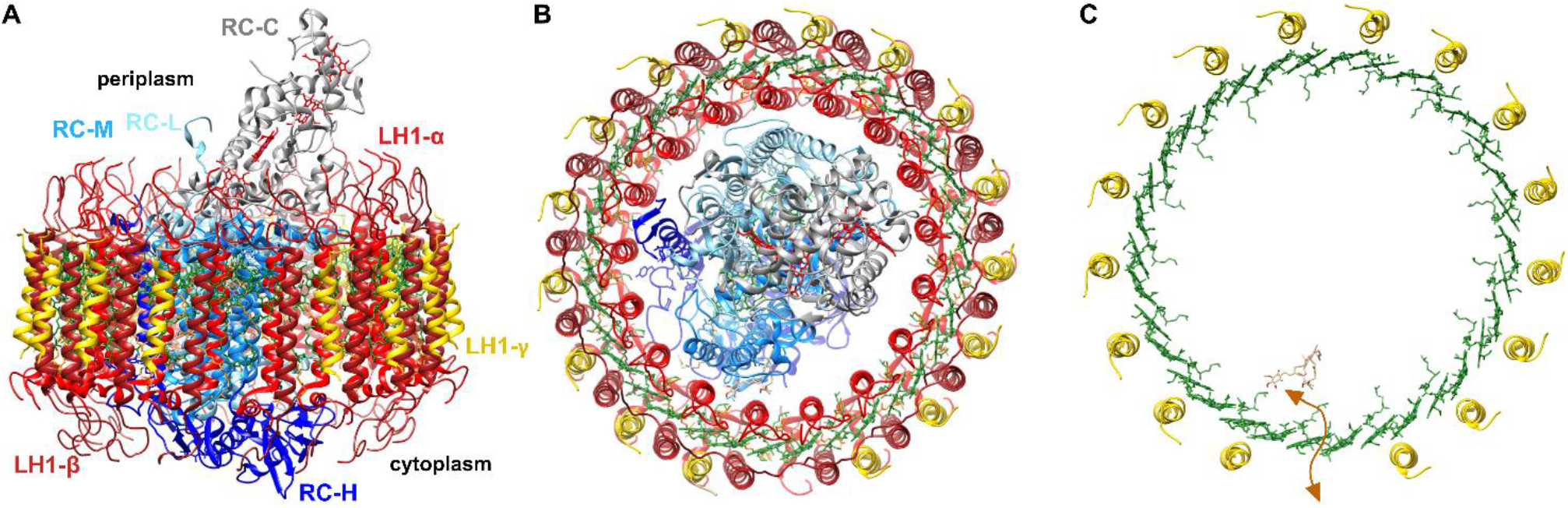
Cryo-EM structure of the *Blc. viridis* RC–LH1 complex, displaying γ subunit locations. A) Side-on view of the RC–LH1 complex, with the periplasmic side facing up. The individual components are indicated in text of the respective colour. The gap created by the ‘missing’ LH1γ polypeptide is at the anterior of the complex. B) View of the RC–LH1 complex from above the periplasmic surface of the membrane, with the gap in LH1 at the bottom of the ring. C) View as in previous with only LH1γ subunits displayed around the LH1 BChls (green). The route for diffusion of quinone/quinol (tan) is indicated by the orange arrow.

Peptide analysis of γ isolated from *Blc. viridis* indicated that the polypeptide is 36 amino acids in length, and variance at position 34 was detected with threonine and valine residues identified in a 2:1 ratio, respectively [22]. This suggested that multiple copies of LH1γ-encoding genes were present in the *Blc. viridis* genome. Subsequent genome sequencing has revealed that *Blc. viridis* has four genes encoding putative γ polypeptides, three of which are clustered and share high sequence identity (LH1γ_1–3_), and the last of which being more divergent (LH1γ_4_) [23,24] (**Fig. 2**).

**Figure 2.**
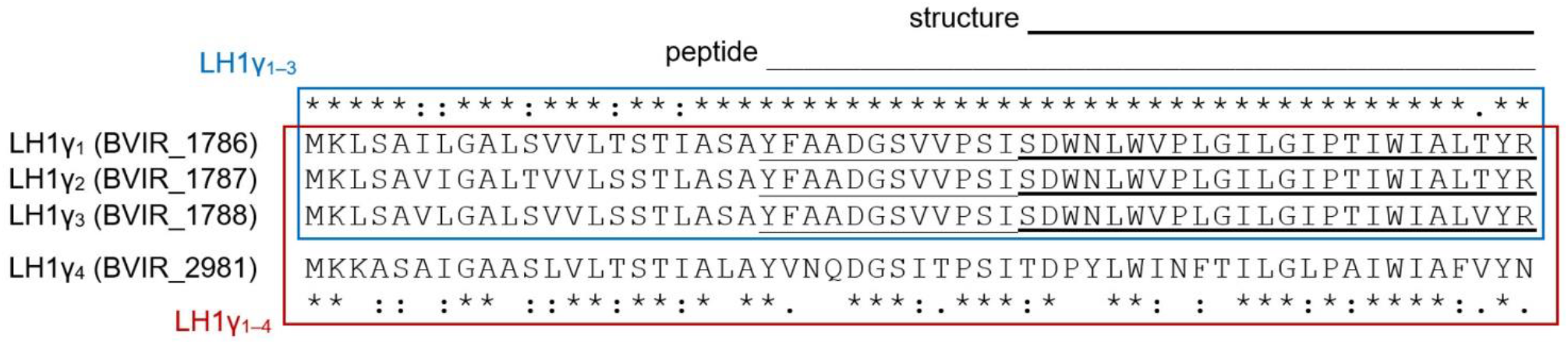
Amino acid sequence alignment of translated LH1γ-encoding genes from *Blc. viridis*. Genetic loci for each ORF are listed in parentheses. Full-length sequence comparisons between clustered LH1γ polypeptides 1–3 (blue box), and with the additional, divergent LH1γ_4_ (red box); identical, highly similar and similar amino acids are indicated by asterisks, colons, and periods, respectively. The sequences of the polypeptides found in the purified complex, identified by Edman degradation [22], and those resolved in the cryo-EM structure [16], are indicated by light and heavy underlining, respectively.

In the present study, we construct mutants of *Blc. viridis* that do not produce LH1γ, permitting the confirmation of the role of this unusual core complex component. Our results indicate that the loss of γ results in large blue-shifts in the absorption maxima of whole cells and of the isolated RC–LH1 complex to a region of the solar spectrum within which photons poorly transmit through the atmosphere, providing an evolutionary rationale for the recruitment of this additional subunit into the bacterial photosystem.

## MATERIALS & METHODS

### Growth of described strains

*Blc. viridis* DSM 133 and mutant strains were grown phototrophically at 30°C in anoxic sodium succinate medium 27 (N27 medium) [26] under illumination (100 μmol·photons·m^−2^·s^−1^) provided by 100W Bellight halogen bulbs or a bank of 24W fluorescent tubes as previously described [27]. All culture transfers were performed in an anoxic chamber (Coy Laboratories). Where required, medium was supplemented with antibiotics at the following concentrations: kanamycin (30 μg·ml^−1^), spectinomycin (30 μg·ml^−1^), carbenicillin (100 μg·ml^−1^). *E. coli* strains JM109 [28] and ST18 (DSM 22074) [29] transformed with pK18mob*sacB* [30] and pBBRBB-derived plasmids [31] were grown in a rotary shaker at 37°C in LB medium supplemented with 30 μg·ml^−1^ kanamycin. ST18 cells were supplemented with 50 μg·ml^−1^ 5-aminolevulinic acid (ALA). All strains and plasmids used in the present study are listed in Supplementary Table 1.

### Construction of deletion mutants of *Blc. viridis*

*Blc. viridis* open reading frames were replaced with genes conferring antibiotic resistance such that their expression is driven by the promoter of the targeted genes, as previously described [32]. Sequences ^~^600 bp up- and downstream of genes encoding LH1γ polypeptides were amplified with relevant UpF and UpR, and DownF and DownR primer pairs, respectively; sequences of the primers used in the present study can be found in Supplementary Table 2. The *aadA* gene from pSRA81 conferring streptomycin/spectinomycin resistance, and the *bla* gene from pET3a conferring resistance to ampicillin/carbenicillin, were amplified from purified plasmids. Resulting amplicons were fused by overlap extension PCR, digested with relevant restriction enzymes, and ligated into similarly digested pK18*mobsacB*. The resulting plasmids were verified by DNA sequencing and conjugated into *Blc. viridis* via *E. coli* ST18. Transconjugants in which the plasmid had integrated into the genome by a single homologous recombination event were selected on N27 medium supplemented with kanamycin. A second recombination event was then promoted by *sacB*-mediated counterselection on N27 medium supplemented with 5% (w/v) sucrose, containing either spectinomycin or carbenicillin, but lacking kanamycin. Sucrose- and spectinomycin/carbenicillin-resistant, kanamycin-sensitive colonies had excised the allelic exchange vector through the second recombination event, and replacement of targeted open reading frames with antibiotic resistance genes was confirmed by colony PCR and sequencing using relevant CheckF and CheckR primers.

### Construction of a replicating expression plasmid for *Blc. viridis*

The *puf* promoter, found upstream of the genes encoding the majority of the LH1 and RC subunits, was amplified from *Blc. viridis* genomic DNA with Ppuf^Bv^F and Ppuf^Bv^R primers. The resulting amplicon was digested with XbaI and BglII, and cloned in place of the *Rhodobacter sphaeroides puf* promoter in pBBRBB-P*puf*_843–1200_-DsRed [31], a broad-host range plasmid containing the pBBR1 origin of replication, generating pBBRBB-*Ppuf^Bv^*.

### Complementation of *Blc. viridis* mutants *in trans*

The genes encoding LH1γ_1_ and LH1γ_4_ were amplified from *Blc. viridis* genomic DNA with the relevant primer pairs, and the resulting amplicons were digested with BglII and SpeI and cloned in place of DsRed-Express2 in pBBRBB-*Ppuf^Bv^*. The resulting plasmids were verified by DNA sequencing and conjugated into the *Blc. viridis* mutant lacking LH1γ from *E. coli* ST18. Transconjugants harbouring the plasmids were selected on N27 medium supplemented with kanamycin, and the presence of the plasmids was confirmed by colony PCR using the primers used to amplify LH1γ_1_ and LH1γ_4_.

### Cell breakage, membrane isolation and solubilisation

Cultures were centrifuged at 4,000 × *g* for 30 min. Cell pellets were resuspended in working buffer (20 mM Tris, 5 mM sodium ascorbate pH 8.0), and were broken by two passages through a cell disruptor (Constant Systems) at 18 000 psi. Unbroken cells were removed by centrifugation at 33 000 × *g* for 15 min. The supernatant was centrifuged at 113,000 x *g* for 2 h in a Beckman Type 45 Ti rotor. Pelleted membranes were resuspended in working buffer and homogenised until no aggregates remained. Resulting membrane suspensions were solubilised by the addition of n-dodecyl β-D-maltoside (β-DDM) to a final concentration of 3% (w/v) with constant gentle stirring for 1 h in the dark, followed by further centrifugation at 150,000 x g for 1 h to remove insoluble material. All steps were carried out at 4°C under dim light.

### Isolation of RC/RC–LH1 complexes

Solubilised membranes were gently layered on top of continuous 15–25% (w/w) sucrose density gradients made up in working buffer containing 0.03% (w/v) β-DDM. Gradients were centrifuged in a Beckman SW41 Ti rotor at 90,000 × *g* for 19 h at 4°C. Pigmented bands containing photosynthetic complexes were collected using a fixed needle and a peristaltic pump.

### Absorption spectroscopy

UV/vis/near-IR absorption spectra were collected on a Cary 3500 spectrophotometer (Agilent Technologies) scanning between 300 and 1100 nm at 1 nm intervals with a 0.1 s integration time.

### Measurement of lightbulb emission

Emission spectra from halogen bulbs and fluorescent tubes were measured on a FlouroMax-4 spectrofluorometer (Horiba). Bulbs were directed into the open sample chamber in a dark room and emission measurements were taken between 365 and 1150 nm at 1 nm intervals with the excitation source switched off.

### Pigment extraction

Bacterial cultures were washed in 50mM Tris-HCl pH 8.0 and pelleted by centrifugation. Total pigments were extracted by adding 10 pellet volumes of 7:2 (v/v) acetone/methanol, immediately vortex-mixed for 30 s, and the resulting suspension incubated on ice for 15 min. Extracts were clarified by centrifugation and the supernatants were filtered through 0.22 μm PVDF membrane filters and immediately analysed. For the preparation of carotenoids, a drop of 5 M NaCl and 10 pellet volumes of hexane were added to the clarified acetone/methanol extract. The sample was mixed and the phases allowed to partition. The upper hexane phase was transferred to a glass vial, dried in a vacuum concentrator at 30°C, and reconstituted in a small volume of acetonitrile immediately prior to analysis by reversed-phase high-performance liquid chromatography (HPLC). Centrifugation steps were performed at 15,000 g for 5 min at 4°C.

### Pigment analysis

Extracted pigments were separated on an Agilent 1100 HPLC system maintained at 40°C with a flow rate of 1 ml·min^−1^. BChls and BPhe pigments were separated on a ThermoFisher Acclaim 120 C18 column (3 μm particle size, 120 Å pore size, 150×3 mm) using a 20 min isocratic gradient of 80:20 (v/v) methanol/acetone. Elution of BChl *b* and bacteriopheophytin *b* (BPhe *b*) was monitored by checking the absorbance at the Soret and Qy absorption maxima of the relevant pigments. Carotenoids were separated on a Supelco Discovery HS C18 column (5 μm particle size, 120 Å pore size, 250×4.6 mm) using a method modified from Magdaong et al. [33]. Pigments were eluted on a 50 min isocratic gradient of 58:35:7 (v/v/v) acetonitrile/methanol/tetrahydrofuran. Elution of carotenoid species was monitored at 470 nm and 505 nm.

## RESULTS

### Deletion of LH1γ-encoding genes prevents absorption of light >1000 nm

In order to determine the role of the LH1γ polypeptides in the *Blc. viridis* RC–LH1, the open reading frames encoding LH1γ_1_, LH1γ_2_, and LH1γ_3_, which are clustered together in the genome, were replaced with a spectinomycin resistance cassette, generating the mutant strain ΔLH1γ_1–3_ (**Fig. 3A**). Subsequently, the more divergent paralog LH1γ_4_ was replaced in the ΔLH1γ_1–3_ background with an ampicillin/carbenicillin resistance cassette, generating strain ΔLH1γ_1–4_ lacking all copies of LH1γ-encoding genes (**Fig. 3B**). The presence of LH1γ_4_ has not been detected in the RC–LH1 of *Blc. viridis*, thus was not removed from the genome of the WT strain.

**Figure 3.**
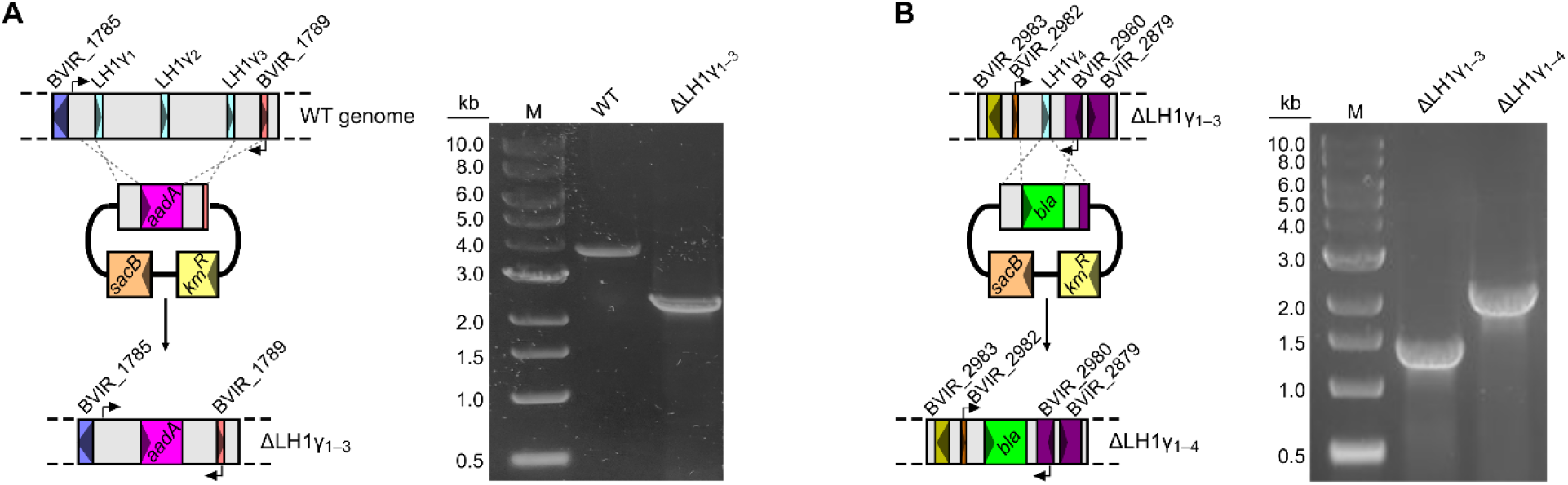
Replacement of LH1γ-encoding genes. Cartoon representations of the arrangement of A) LH1γ_1–3_, and B) LH1γ_4_-encoding genes (left), and their replacement with antibiotic resistance cassettes. Arrows represent binding sites of primers used for screening via colony PCR, and resulting agarose gels confirming replacement are shown (right).

BChl-dependent anoxygenic phototrophs have classically been cultured with illumination provided by incandescent bulbs, which emit a broad spectrum of light from the visible deep into the infrared [34]. Since incandescent light bulbs have been banned in many parts of the world (e.g. phased out in the European Union by 2012), many researchers working on these organisms switched to halogen bulbs [35,36], which have now also been banned for production but can still be sourced. Therefore, our standard laboratory conditions for culturing anoxygenic phototrophic proteobacteria is illumination at 100 μmol·photons·m^−2^·s^−1^ provided by halogen bulbs (see **Fig. S1** for bulb emission spectra). The ΔLH1γ_1–3_ and ΔLH1γ_1–4_ mutants, along with the WT, were cultured under these conditions, and their whole-cell absorption spectra were measured (**Fig. 4**). The WT displayed the characteristic absorption band with a maximum at 1018 nm. Interestingly, the spectra obtained for the ΔLH1γ_1–3_ and ΔLH1γ_1–4_ strains displayed a prominent absorption peak at 825 nm, and a small absorption feature at approximately 970 nm. This absorption profile is similar to that of the purified RC from *Blc. viridis* [37], and to membranes isolated from a mutant of BChl *a*-synthesising *Rhodobacter sphaeroides* containing RCs but lacking any antenna complexes, albeit with absorption maxima blue-shifted from those of the BChl *b*-containing mutants described here [38]. In the context of these studies, our results indicate that mutants ΔLH1γ_1–3_ and ΔLH1γ_1–4_ are unable to assemble LH1 in the absence of LH1γ polypeptides, when grown under our standard laboratory conditions. This suggested that loss of LH1γ may severely affect the stability of the LH1 ring, or prevents its assembly altogether.

**Figure 4.**
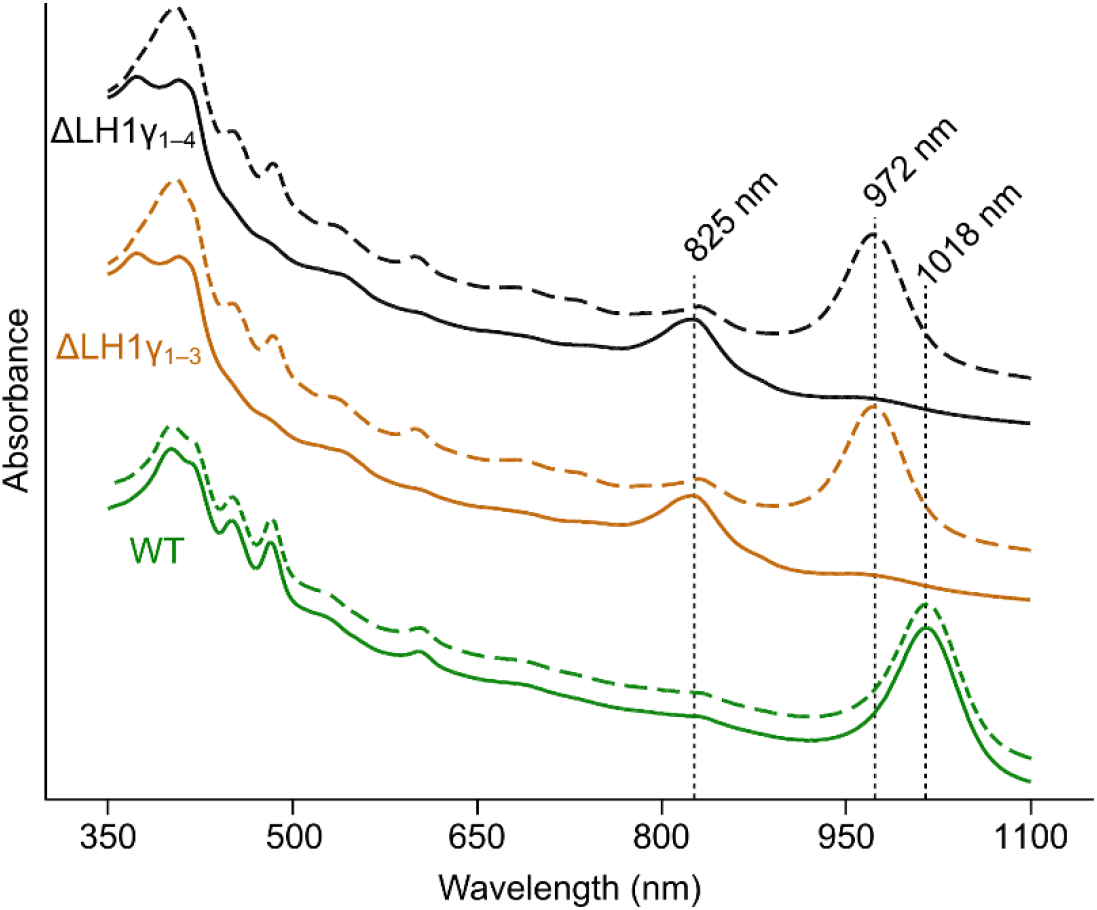
Whole-cell absorption spectra of *Blc. viridis* strains. Spectra of cultures of WT (green traces), ΔLH1γ_1–3_ (orange traces), and ΔLH1γ_1–4_ (black traces), grown under halogen light (solid lines) or white light (dashed lines) were recorded. Major absorption bands in the near-IR region of the spectrum are indicated.

The absorption features in the 400–550 nm range of the whole-cell spectra of the ΔLH1γ_1–3_ and ΔLH1γ_1–4_ mutants grown under halogen light indicate that the cells accumulate carotenoids, accessory pigments that absorb in this range and also play roles in pigment–protein complex stability and quenching of triplet excited states of (B)Chls [39]. The ΔLH1γ_1–3_ and ΔLH1γ_1–4_ mutants, along with the WT, were cultured under white light from fluorescent tubes in order to provide carotenoid-specific wavelengths that would not be captured by the BChls in the RC. Under these conditions the strains were phototrophically viable and the whole-cell absorption spectra were measured (**Fig. 4**). As under halogen light, the WT displayed characteristic absorption with a maximum at 1018 nm. Surprisingly, ΔLH1γ_1–3_ and ΔLH1γ_1–4_ both displayed spectra with absorption maxima at 972 nm, 46 nm blue-shifted with respect to the WT maximum. These mutant spectra displayed an absorption profile more similar to the WT and unlike those of the RC alone, suggesting that these strains assemble the LH1 ring around the RC, albeit without LH1γ. Interestingly, strain ΔLH1γ_1–3_, which still contains LH1γ_4_, displays the same absorption spectra as the mutant lacking all encoding genes; the ΔLH1γ_1–4_ strain completely lacking LH1γ was subsequently used for all experiments.

### LH1 assembles around the RC without LH1γ only under white light

In order to analyse the pigment–protein complexes assembled in the described strains, they were cultured under halogen light and white light, the membranes from these cells were solubilised with detergent, and the resulting membrane complexes were subjected to rate-zonal centrifugation on continuous sucrose density gradients (**Fig. 5**). The WT grown under both halogen light and white light accumulates a sole pigmented complex, each displaying identical densities and absorption spectra. The mutant lacking LH1γ grown under halogen light accumulates a single complex at greatly reduced density compared to that of the WT; the absorption spectrum of this band displays the characteristic profile of an isolated BChl *b*-containing RC. When this mutant is grown under white light, RCs can also be identified in the sucrose gradients, but a second pigmented complex, with a density slightly less than that of the WT RC–LH1, is also present as the major band. The absorption spectrum of this band displays a maximum at 965 nm, slightly blue shifted from its whole-cell absorbance maximum, as is common for isolated complexes.

**Figure 5.**
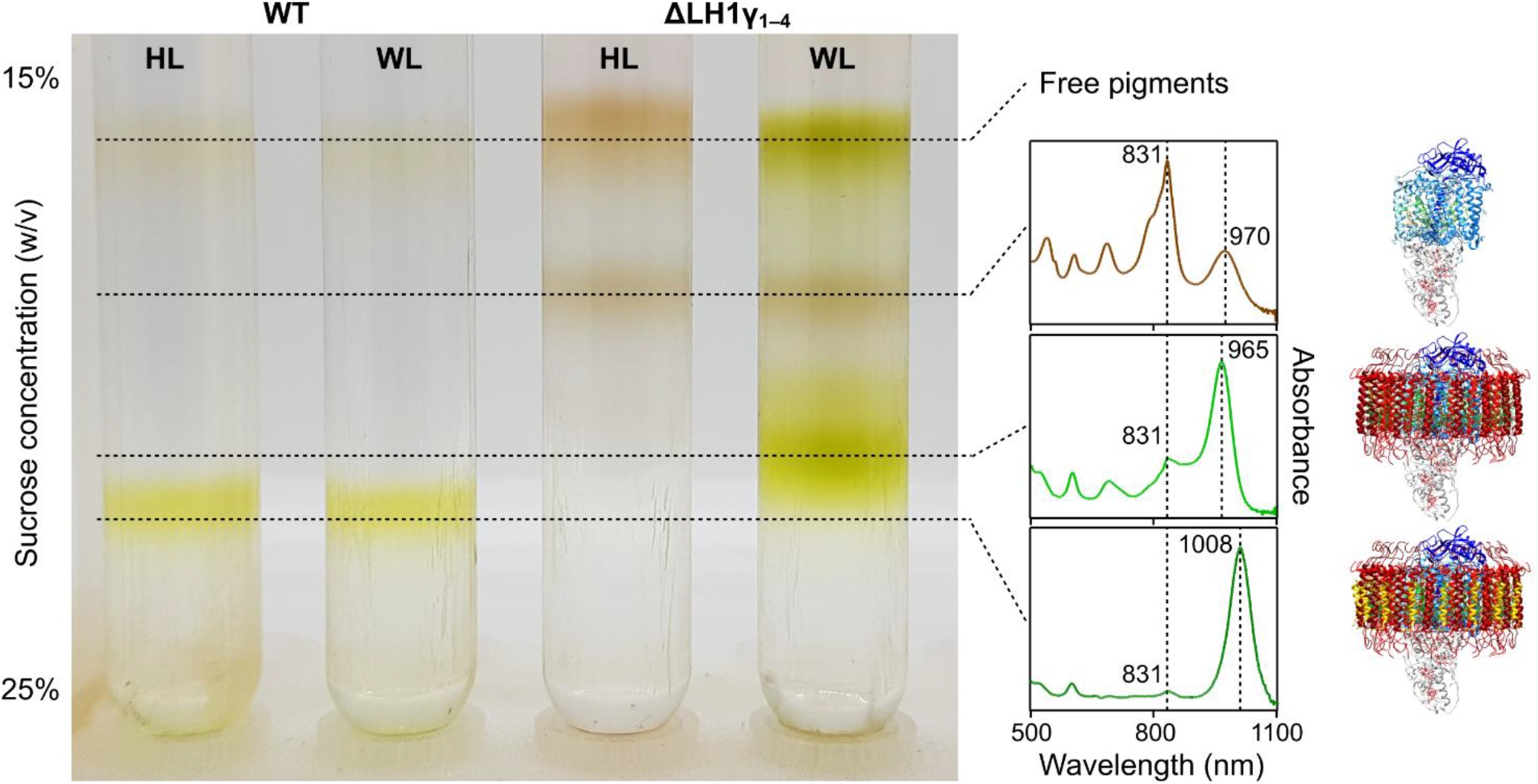
Sucrose density gradients of solubilised membranes from WT and LH1γ-lacking strains of *Blc. viridis*. Absorption spectra of relevant isolated bands are indicated by dashed lines, and the presumed structures of these complexes are shown to the right of their respective spectra for illustrative purposes. HL, halogen light; WL, white light.

The major pigmented band from each of the gradients was isolated and buffer-exchanged in a spin concentrator to remove sucrose. The complexes were denatured in SDS sample buffer and the subunits separated via Tris–tricine gel electrophoresis (**Fig. S2**). The stained gel indicates that the major complex from the LH1γ-lacking mutant grown under halogen light does not contain any LH1 components, including LH1β and LH1α. However, the major complex isolated from this strain grown under white light contains LH1β and LH1α, but as expected lacks LH1γ, confirming that this blue-shifted complex is an intact RC–LH1.

### Loss of LH1γ slows the phototrophic growth rate under halogen light

To assess the effect of deletion of the genes encoding LH1γ, the WT and ΔLH1γ_1–4_ strains were cultured under white light conditions to mid exponential phase, standardised by OD_700_ and used to inoculate biological triplicates in fresh medium incubated under halogen light and white light. When grown under halogen light, the doubling time of the mutant was almost double that of the WT (**Table 1**). However, the growth rates measured under white light, when both strains assemble intact RC–LH1, were comparable, albeit much slower than under halogen light, and each strain displaying inconsistent growth across the triplicate cultures.

**Table 1.** Phototrophic growth rates of *Blc. viridis* strains under different light regimes. Doubling times were calculated from biological triplicates under each condition.

| Strain | Doubling time (h) |  |
| --- | --- | --- |
|  | Halogen light | White light |
| WT | 12.8 ± 0.38 | 94.8 ± 18.3 |
| ΔLH1$\gamma_{1-4}$ | 25.3 ± 0.63 | 90.1 ± 20.4 |

### Pigment analysis of the LH1γ mutant

The RCs of purple bacteria assemble with four BChls, two bacteriopheophytin molecules (demetallated analogues of their parent BChl) and a ‘kinked’ *cis*-carotenoid [2,40]. The 17 LH1 subunits of the *Blc. viridis* core complex (16 αβγ trimers and an αβ pair) each coordinate two BChl *b* pigments and a linear ‘all-*trans*’ carotenoid [16], meaning that the total pigment content of its RC–LH1 is 38 BChl *b*, 2 BPhe *b*, and 18 carotenoids (**Fig. S3**). The RC of *Blc. viridis* is known to contain 15-*cis*-1,2-dihydroneurosporene, but the cells also accumulate a range of neurosporene- and lycopene-type carotenoids [10,41]. To analyse the pigment composition of WT and ΔLH1γ_1–4_ cultured under both light regimes, biological triplicates of strains grown to late exponential phase were collected, standardised by cell number, and pellets of these were subjected to pigment extraction with an excess of organic solvent. The standardised extracts were subjected to analysis by HPLC. Representative traces, normalised to major peak height for clarity, demonstrating separation of BChl *b* and BPhe *b* are shown in **Fig. 6A**, and those demonstrating the separation of carotenoid species are shown in **Fig. 6B. Fig. 6C** shows the contents of each pigment relative to those found in the WT under our standard growth conditions, hereafter the ‘control’ sample.

**Figure 6.**
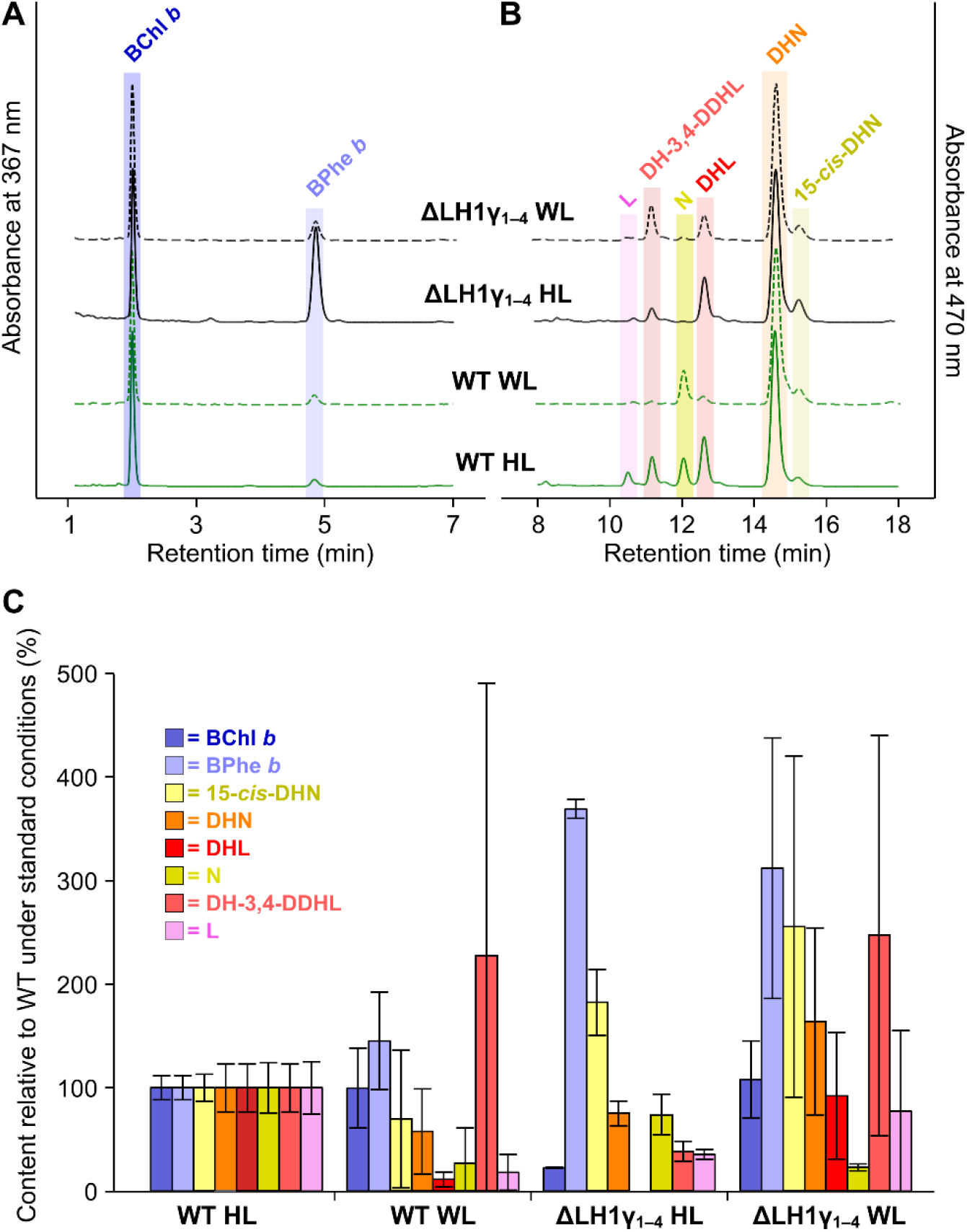
HPLC analysis of pigments accumulated by described strains under halogen and white light. Representative traces of **A)** BChl and BPhe separation, monitored at 367 nm, and **B)** carotenoid species separation, monitored at 470 nm. Traces are normalised to major peak height for clarity. **C)** The cellular content of each pigment from three biological replicates, relative to those found in the WT grown under standard laboratory conditions (100 μmol photons m^−2^ s^−1^ provided by halogen bulbs). Standard deviations for each pigment are shown by the error bars. 15-*cis*-DHN; 15-*cis*-1,2-dihydroneurosporene, DHN, 1,2-dihydroneurosporene; DHL; 1,2-dihydrolycopene, N; neurosporene, DH-3,4-DDHL, 1,2-dihydro-3,4-didehydrolycopene, L; lycopene.

When grown under white light, the WT accumulates a very similar amount of BChl *b* to when grown under halogen light (1.3% less), but 45.4% more BPhe *b* (**Fig. 6C**); the lack of an RC band in the sucrose density gradient from this condition (see **Fig. 5**) indicates that there may be greater turnover of BChl *b* when the cells are grown under white light. When the ΔLH1γ_1–4_ strain is grown under halogen light it displays a considerably higher BPhe:BChl ratio than when grown under white light, and when compared to the WT grown under either condition (**Fig. 6A,C**), this equates to 22.9% of the BChl *b* content and 369.3% of the BPhe *b* content of the control (**Fig. 6C**). This is unsurprising since the mutant grown under halogen light solely assembles RCs without LH1 (see **Fig. 5**). When grown under white light, ΔLH1γ_1–4_ accumulates 8.2% more BChl *b* but more than three times the BPhe *b* content of the control, and 9% more BChl *b* and 2.15 times more BPhe *b* than the WT under white light; despite primarily assembling RC–LH1 under this condition, some RCs lacking the antenna are found in the sucrose gradient (see **Fig. 5**), which may explain this increase in the BPhe *b* content.

The contents of each carotenoid species differ considerably in both strains from those in the control. Using 15-*cis*-1,2-dihydroneurosporene as a marker for the RC, the mutant displays increases of 82.4% and 155.6% in the content of this carotenoid under halogen and white light, respectively, compared to the control, which correlates with the presence of RCs in their sucrose gradients (see **Fig. 5**). The WT grown under white light accumulates 29.7% less of this carotenoid than when grown under halogen light, despite displaying a large amount of variance. This implies that the overall cellular RC–LH1 content may be lower than the control, perhaps as a result of a smaller surface area of internal membrane in which to house it. The carotenoid 1,2-dihydro-3,4-didehydrolycopene accumulates in both strains when grown under white light, the WT displaying 227.4% of the control content, and the mutant having 2.5 times that of the control and 6.4 times more than when it is grown under halogen light. The data shown in **Fig. 6C** indicates that the synthesis of *Blc. viridis* carotenoids, and perhaps the regulation of this process, is dependent on the source of light used for growth. However, the slow doubling times and large variations displayed by both WT and mutant grown under white light may also play a role in the complicated carotenoid profiles of these strains.

### Complementation of the mutant lacking LH1γ *in trans* restores absorption past 1000 nm

To determine if the addition of native *Blc. viridis* genes encoding LH1γ are able to restore the *in vivo* absorption maximum of the ΔLH1γ_1–4_ mutant to that of the WT, genes encoding LH1γ_1_ (which is identical to LH1γ_2_, and differs by a single amino acid to LH1γ_3_) and LH1γ_4_, which is more divergent (see **Fig. 2**), were cloned into the purple bacterial expression vector pBBRBB-P*puf*_843–1200_ [31], in place of DsRed. These constructs were conjugated into ΔLH1γ_1–4_, and the transconjugants were cultured under halogen light and white light. The whole-cell absorption spectra of these strains were unchanged from that of the untransformed mutant strain (data not shown). These plasmids contain the strong *puf* promoter from the Rhodobacterterales member *Rhodobacter sphaeroides*, which may not drive expression in a distantly-related member of the Rhizobiales such as *Blc. viridis* [42]. To overcome this, the *Rhodobacter sphaeroides* promoter was replaced with the native *puf* promoter, which has been shown to drive expression of genes in *Blc. viridis* from a replicating plasmid [43]. These constructs were conjugated into ΔLH1γ_1–4_, and the whole cell spectra of these strains grown under halogen light were recorded (**Fig. 7**). The transconjugant harbouring the plasmid-borne gene encoding LH1γ_1_ displayed a whole-cell absorption maximum at 1018 nm, identical to that of the WT. Interestingly, the strain expressing LH1γ_4_ displayed an absorption maximum red-shifted from that of mutant lacking LH1γ, absorbing maximally at 1003 nm, but this maximum is 15 nm shorter than that of the WT or the mutant complemented with LH1γ_1_. This indicates that the amino acid sequence of LH1γ influences the interaction with the neighbouring LH1β polypeptides and determines the extent of the red-shift in the absorption profile of RC–LH1.

**Figure 7.**
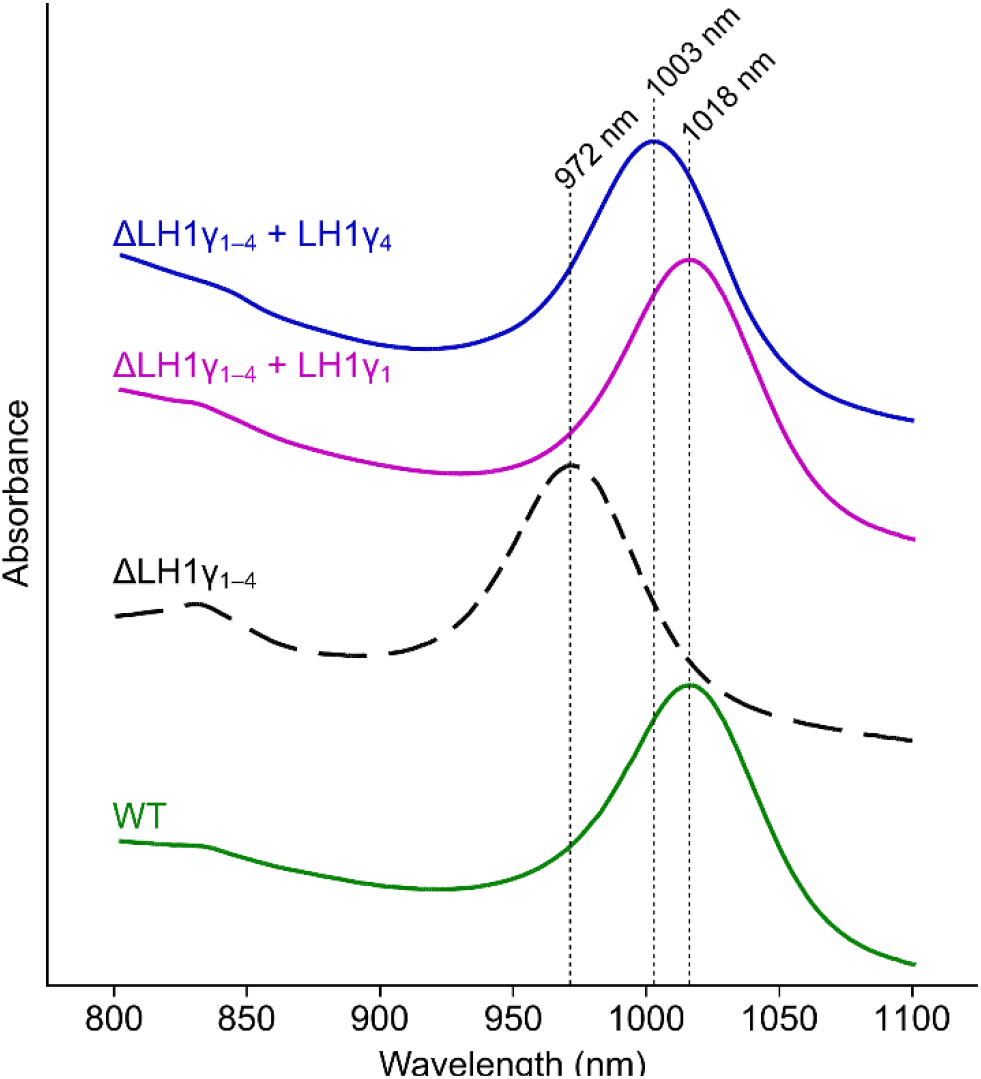
Whole-cell absorption spectra of complemented *Blc. viridis* strains. Spectra of cultures of WT (green trace), ΔLH1γ_1–4_ (black trace), ΔLH1γ_1–4_ + pBBRBB-P*puf*^Bv^ [LH1 γ_1_] (pink trace), and ΔLH1γ_1–4_ + pBBRBB-P*puf^Bv^* [LH1 γ_4_] (blue trace). Cultures were grown under halogen light (solid lines) or white light (dashed line). Major absorption bands in the near-IR region of the spectrum are indicated.

## DISCUSSION

The vast majority of anoxygenic phototrophs in diverse bacteria phyla discovered to date use BChl *a* as the RC pigment for phototrophy [44]. Only a small number of purple bacteria use BChl *b* [4]. Of these, only *Blastochloris* spp. have genes encoding LH1γ. In each case, the common components of the photosystem are encoded by single ORFs within the photosynthesis gene cluster (PGC), which also contains genes encoding pigment biosynthesis enzymes and photosystem assembly factors [45]. However, each sequenced strain contains multiple LH1γ genes that are located distant from the PGC [24,25,46,47]. This suggests that these genes evolved separately from the other PGC genes, and that assembly of RC–LH1 without LH1γ is safeguarded against by the presence of multiple paralogous genes in the genome. The evolutionary rationale for this is provided by our results; our mutant strains assembling RC–LH1 lacking LH1γ absorb maximally at 972 nm, which sits in a range of the solar spectrum that poorly transmits through the atmosphere due to absorption by water vapour (**Fig. 8**). In this range, few photons reach the surface of Earth, and those that do also poorly transmit through liquid water at the surface [48]. Therefore, it is likely that *Blastochloris* spp. evolved or recruited an additional LH1 subunit to influence absorption of the complex, shifting to a region of the spectrum where photons are of lower energy but are more abundant.

**Figure 8.**
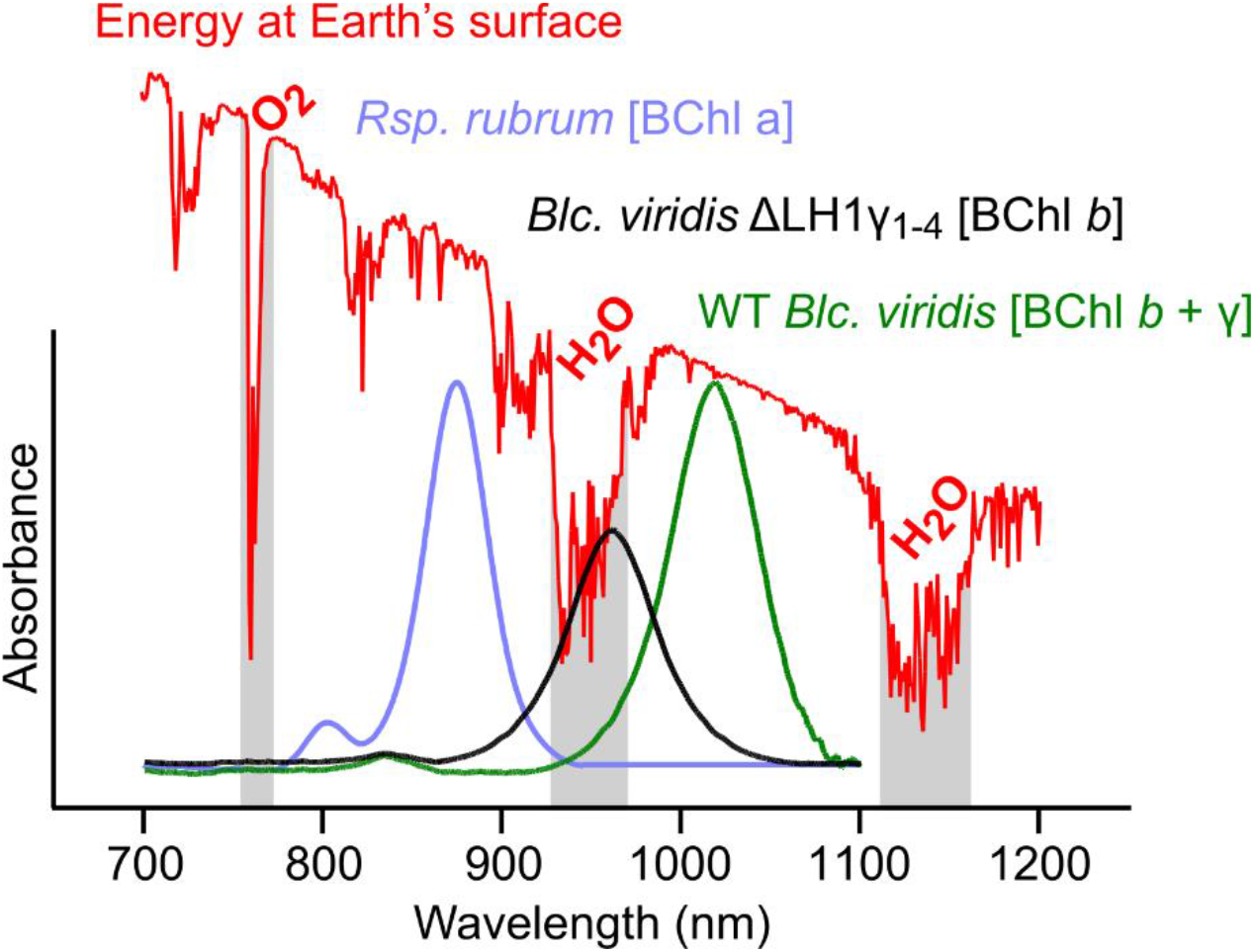
Absorption spectra of RC–LH1 complexes relative to available solar energy. The purified complex from *Rsp. rubrum* containing BChl *a* (blue trace) is blue-shifted in comparison the BChl *b*-containing complexes from WT *Blc. viridis* (green trace) and the ΔLH1γ_1–4_ mutant (black trace). The overlayed red trace displays the spectrum of solar radiation that reaches the surface of the Earth [49]. Absorption bands from O_2_ and H_2_O in the atmosphere are labelled, and the corresponding spectral regions from which photons poorly transmit are highlighted in grey.

LH1γ polypeptides 1 and 2 are identical, and LH1γ_3_ differs by a single amino acid (see **Fig. 2**); their encoding genes are clustered in the genome. The original proteomic analysis of *Blc. viridis* demonstrated that highly similar polypeptides containing threonine or valine at position 34 could be isolated from its photosystem and are present at a 2:1 ratio, while a polypeptide equivalent to LH1γ_4_ was not detected [22]. This indicates that the genes encoding LH1γ_1–3_ are transcribed together, and that LH1γ_4_ is not produced when the cells are grown under full-spectrum light from incandescent bulbs. Similarly, we could not detect any phenotypic change in the cells of the ΔLH1γ_1–3_ mutant when compared to the mutant lacking all γ-encoding genes under any condition tested that we could ascribe to the production of LH1γ_4_. Our finding that expression of the LH1γ_4_ gene from a plasmid in the ΔLH1γ_1–4_ mutant leads to the cells having an absorption maximum of 1003 nm indicates that incorporation of γ_4_ into RC–LH1 induces a moderate red-shift, which may provide more favourable absorption under certain environmental conditions (e.g. when in competition with other BChl *b* phototrophs such as *Halorodospira* spp. displaying similar absorption at 1018 nm, see below). It may be that incorporation of LH1γ_4_ into the WT RC–LH1 can be induced by growth under specific LED regimes in the laboratory, which we intend to explore further.

We found that the ΔLH1γ_1–3_ and ΔLH1γ_1–4_ mutants grown under our standard laboratory conditions, with illumination provided by halogen bulbs, resulted in the assembly of RCs, but no intact RC–LH1 complexes. However, growth under white-light fluorescent tubes that would be routinely used for cultivation of oxygenic cyanobacteria permitted assembly of RC–LH1, with an absorption maximum far outside the range of emission from the light source. This appears counterintuitive, but it is possible that the emission from the halogen bulbs, which reduces sharply at around 850 nm (see **Fig. S1**), provides wavelengths that efficiently excite the BChls contributing the major absorption feature of the RC at 831 nm (see RC band spectrum in **Fig. 5**). This RC contains only a single 15-*cis*-1,2-dihydroneurosporene carotenoid, providing limited absorption in the visible range of the spectrum. The observation that RC–LH1 lacking γ only assemble when the mutants are grown under white light might be explained by the carotenoid content of LH1; the 17 all-*trans* pigments bound in ring provide strong absorption in the 400–550 nm range, which overlaps with the narrow, intense emission band of the fluorescent tubes in this range (see **Fig. S1**). Our results will be noteworthy to those studying anoxygenic phototrophy; although it is possible to purchase LED panels that mimic ‘photosynthetically-active radiation’ for oxygenic phototrophs, panels emitting wavelengths to ^~^1200 nm that mimic full-spectrum natural sunlight do not yet exist. As it becomes more difficult to source incandescent and halogen bulbs there may be a need for researchers to assemble broad-spectrum LED regimes themselves, to accurately study phototrophic bacteria under conditions similar to those experienced in nature.

The mutants lacking LH1γ assembling RC–LH1 in white light also accumulate RCs, as can be seen in the sucrose density gradients, while the WT does not (**Fig. 5**). This suggests that the presence of the 16 LH1γ subunits provide a stabilising role; the interaction with the adjacent β polypeptides may fix the complex in place in addition to imparting the red-shift in absorption, and the presence of γ may orient the RC in such a position that efficient quinone/quinol shuttling can occur at the ‘missing’ 17^th^ γ position [16]. This stabilising effect may be analogous to the role that PufX plays in the primarily dimeric *Rhodobacter sphaeroides* RC–LH1 [50]. Deletion of the *pufX* gene leads to formation of solely monomeric RC–LH1, and the recent cryo-EM structure of this complex was solved at much lower resolution than the PufX-containing dimer or monomer, attributed to PufX’s role in orienting the RC within LH1 [50,18]. It is interesting that we do not observe assembly of RCs without LH1 in the WT under either white light or halogen light conditions. This could be due to the stability imparted by γ; it is likely that the formation of RC–LH1 is a highly coordinated process and that γ is incorporated into LH1 at the same time as the α and β polypeptides by the assembly machinery, and this may not be easily reversed once γ has stabilised the complex. Turnover and recycling of light-harvesting and RC complexes is a well-studied process; photosystem II of oxygenic phototrophs is damaged by the reaction in catalyses so is constantly reassembled [51], and the metabolically diverse purple bacteria using BChl *a* can assemble and disassemble RC–LH1 depending on whether conditions suit aerobic respiration [52]. However, *Blastochloris* spp. are unable to grow in the presence of O_2_ and they generally inhabit anoxic ecological niches such as the bottom of microbial mats, where rapid turnover of the photosystem is not required [53]. Here, extremely red-shifted absorption provided by the γ-stabilised complex affords a competitive advantage over the abundant BChl *a*-containing phototrophs in the mat where wavelengths are highly filtered, and thus a route to survival and growth.

*Halorhodospira (Hrs.) halochloris* and *Hrs*. *abdelmalekii* (formerly *Ectothiorhodospira halochloris* and *Ectothiorhodospira abdelmalekii*) are additional BChl *b*-containing purple bacteria isolated from hypersaline and alkaline soda lakes in Wadi El Naturn, Egypt, that also display whole-cell absorption maxima at 1018 nm [54,55]. The isolated RC–LH1 complexes from these organisms were found to contain three low molecular weight components, and the LH1 absorption at 1018 nm could be reversibly blue-shifted to 964 nm when titrated with acid in the pH range of 7.5–5.7 [56]. The small polypeptide of a similar size to the *Blc. viridis* LH1γ was isolated from the RC–LH1 complex of *Hrs. halochloris* and was assigned to be analogous, although its sequence was not presented in the publication [57]. The recent complete genome sequence of *Hrs. halochloris* has revealed than an ortholog of a *Blastochloris* spp. gene encoding LH1γ is absent [58]. The inability to find candidates encoding LH1γ in the genome of *Hrs. halochloris—as* well as in other non-*Blastochloris* BChl *b*-containing organisms—via BLAST search is unsurprising since the polypeptide is a small, single transmembrane helix (e.g. the sequences of the single transmembrane PufX polypeptides from *Rhodobacter* spp. display low sequence identity across the genus [59]). *Hrs. halochloris* was found to have an unusual complement of genes encoding RC–LH1 subunits [58]. This organism has 2 *pufBALMC* operons, and an additional 2 pairs of *pufBA* genes found elsewhere in the genome, one pair of which (*pufB3A3*) is more divergent, sharing only 41% and 42% identity to LH1β1 and LH1α1, respectively. Recent results demonstrate that desalting the purified *Hrs. halochloris* RC–LH1 complex results in the same blue-shift described previously with a decrease in pH, which can be reversed by supplementation with salts [60]. Taking all of these results together, it is likely that *Halorhodospira* spp. have evolved an alternative approach to accessing photons of wavelengths >1000 nm to that taken by members of the *Blastochloris* genus; the polypeptide thought to be analogous to LH1γ is likely LH1β or LH1α encoded by a paralogous ORF in the *Hrs. halochloris* genome, resulting in LH1 containing multiple forms of α- and/or β-polypeptides, as were found in the RC–LH1 of BChl *a*-containing *Thiorhodovibrio* strain 970 [61].

This study elucidates the role of the γ polypeptide in the RC–LH1 of *Blc. viridis*. The presence of the highly conserved γ isoform stabilises the complex and shifts its absorption 46 nm further into the near-infrared. This extreme red-shift allows *Blastochloris* spp. to access the more abundant photons >1000 nm that are not absorbed by water in the atmosphere. The discovery that plasmid-borne expression of a divergent isoform of LH1γ not found in the WT complex results in only a moderate red-shift of the complex upon assembly provides a route to explore the extent to which polypeptide sequence influences the absorption shift, and whether the low energy limit of natural photosynthesis can be surpassed by rational design of the LH1γ subunit.

## AUTHOR CONTRIBUTIONS

DPC conceived the study and designed the experiments with significant input from DN and NMR. All authors conducted experiments and analysed the results. DPC wrote the manuscript with contributions from DN and NMR.

## ACKNOWLEDGEMENTS

This work was supported by grants from the Royal Society [RGS\R2\192311], the Wellcome Trust [ISSF 204822/Z/16/Z], and University of Liverpool start-up funds to DPC. DN is supported by a Mahidol–Liverpool PhD Scholarship. NMR is supported by a Doctoral Training Partnership from the Biotechnology and Biological Sciences Research Council (BBSRC). DPC acknowledges support from BBSRC grant BB/W008076/1.

## REFERENCES

1. Bryant DA, Canniffe DP. How nature designs light-harvesting antenna systems: design principles and functional realization in chlorophototrophic prokaryotes. Journal of Physics B: Atomic, Molecular and Optical Physics. 2018 Jan 17;51(3):033001.

2. Blankenship RE. Molecular mechanisms of photosynthesis. John Wiley & Sons; 2021 Jul 20.

3. Partensky F, Hess WR, Vaulot D. Prochlorococcus, a marine photosynthetic prokaryote of global significance. Microbiology and molecular biology reviews. 1999 Mar 1;63(1):106–27.

4. Hoogewerf GJ, Jung DO, Madigan MT. Evidence for limited species diversity of bacteriochlorophyll b-containing purple nonsulfur anoxygenic phototrophs in freshwater habitats. FEMS microbiology letters. 2003 Jan 1;218(2):359–64.

5. Eimhjellen KE, Aasmundrud O, Jensen A. A new bacterial chlorophyll. Biochemical and Biophysical Research Communications. 1963 Feb 6;10(3):232–6.

6. Drews G, Giesbrecht P. Die Thylakoidstrukturen von Rhodopseudomonas spec. Archiv für Mikrobiologie. 1965 Sep;52(3):242–50.

7. Drews G, Giesbrecht P. Rhodopseudomonas viridis, nov. spec., ein neu isoliertes, obligat phototrophes Bakterium. Archiv für Mikrobiologie. 1966 Sep;53(3):255–62.

8. Hiraishi A. Transfer of the Bacteriochlorophyll b-Containing Phototrophic Bacteria Rhodopseudomonas viridis and Rhodopseudomonas sulfoviridis to the Genus Blastochloris gen. nov. International Journal of Systematic and Evolutionary Microbiology. 1997 Jan 1;47(1):217–9.

9. Deisenhofer J, Epp O, Miki K, Huber R, Michel H. X-ray structure analysis of a membrane protein complex: electron density map at 3 Å resolution and a model of the chromophores of the photosynthetic reaction center from Rhodopseudomonas viridis. Journal of molecular biology. 1984 Dec 5;180(2):385–98.

10. Deisenhofer J, Epp O, Miki K, Huber R, Michel H. Structure of the protein subunits in the photosynthetic reaction centre of Rhodopseudomonas viridis at 3Å resolution. Nature. 1985 Dec;318(6047):618–24.

11. Roszak AW, Howard TD, Southall J, Gardiner AT, Law CJ, Isaacs NW, Cogdell RJ. Crystal structure of the RC-LH1 core complex from Rhodopseudomonas palustris. Science. 2003 Dec 12;302(5652):1969–72.

12. Jackson PJ, Hitchcock A, Swainsbury DJ, Qian P, Martin EC, Farmer DA, Dickman MJ, Canniffe DP, Hunter CN. Identification of protein W, the elusive sixth subunit of the Rhodopseudomonas palustris reaction center-light harvesting 1 core complex. Biochimica et Biophysica Acta (BBA)-Bioenergetics. 2018 Feb 1;1859(2):119–28.

13. Lilburn TG, Haith CE, Prince RC, Beatty JT. Pleiotropic effects of pufX gene deletion on the structure and function of the photosynthetic apparatus of Rhodobacter capsulatus. Biochimica et Biophysica Acta (BBA)-Bioenergetics. 1992 May 20;1100(2):160–70.

14. Farchaus JW, Barz WP, Grünberg H, Oesterhelt D. Studies on the expression of the pufX polypeptide and its requirement for photoheterotrophic growth in Rhodobacter sphaeroides. The EMBO journal. 1992 Aug;11(8):2779–88.

15. Bracun L, Yamagata A, Christianson BM, Terada T, Canniffe DP, Shirouzu M, Liu LN. Cryo-EM structure of the photosynthetic RC-LH1-PufX supercomplex at 2.8-Å resolution. Science Advances. 2021 Jun 1;7(25):eabf8864.

16. Qian P, Swainsbury DJ, Croll TI, Salisbury JH, Martin EC, Jackson PJ, Hitchcock A, Castro-Hartmann P, Sader K, Hunter CN. Cryo-EM structure of the monomeric Rhodobacter sphaeroides RC–LH1 core complex at 2.5 Å. Biochemical Journal. 2021 Oct 29;478(20):3775–90.

17. Tani K, Nagashima KV, Kanno R, Kawamura S, Kikuchi R, Hall M, Yu LJ, Kimura Y, Madigan MT, Mizoguchi A, Humbel BM. A previously unrecognized membrane protein in the Rhodobacter sphaeroides LH1-RC photocomplex. Nature communications. 2021 Nov 2;12(1):1–9.

18. Cao P, Bracun L, Yamagata A, Christianson BM, Negami T, Zou B, Terada T, Canniffe DP, Shirouzu M, Li M, Liu LN. Structural basis for the assembly and quinone transport mechanisms of the dimeric photosynthetic RC–LH1 supercomplex. Nature communications. 2022 Apr 13;13(1):1–2.

19. Barz WP, Vermeglio A, Francia F, Venturoli G, Melandri BA, Oesterhelt D. Role of the PufX protein in photosynthetic growth of Rhodobacter sphaeroides. 2. PufX is required for efficient ubiquinone/ubiquinol exchange between the reaction center QB site and the cytochrome bc1 complex. Biochemistry. 1995 Nov 1;34(46):15248–58.

20. Swainsbury DJ, Qian P, Jackson PJ, Faries KM, Niedzwiedzki DM, Martin EC, Farmer DA, Malone LA, Thompson RF, Ranson NA, Canniffe DP. Structures of Rhodopseudomonas palustris RC-LH1 complexes with open or closed quinone channels. Science Advances. 2021 Jan;7(3):eabe2631.

21. Jay F, Lambillotte M, Mühlethaler K. Localisation of Rhodopseudomonas viridis reaction centre and light harvesting proteins using ferritin-antibody labelling. European journal of cell biology. 1983 Mar 1;30(1):1–8.

22. Brunisholz RA, Jay F, Suter F, Zuber H. The light-harvesting polypeptides of *Rhodopseudomonas viridis*. The complete amino-acid sequences of B1015-alpha, B1015-beta and B1015-gamma. Biological Chemistry Hoppe-Seyler. 1985 366(1):87–98.

23. Qian P, Siebert CA, Wang P, Canniffe DP, Hunter CN. Cryo-EM structure of the *Blastochloris viridis* LH1–RC complex at 2.9 Å. Nature. 2018 556(7700):203–8.

24. Tsukatani Y, Hirose Y, Harada J, Misawa N, Mori K, Inoue K, Tamiaki H. Complete genome sequence of the bacteriochlorophyll b-producing photosynthetic bacterium Blastochloris viridis. Genome Announcements. 2015 Sep 3;3(5):e01006–15.

25. Liu LN, Faulkner M, Liu X, Huang F, Darby AC, Hall N. Revised genome sequence of the purple photosynthetic bacterium Blastochloris viridis. Genome announcements. 2016 Jan 21;4(1):e01520–15.

26. Claus D., Schaub-Engels C. (Eds.), German Collection of Microorganisms: Catalogue of Strains (2nd ed.), DSMZ Catalogue of Microorganisms, Braunschweig, Germany (1977), pp. 279–280

27. Canniffe DP, Hunter CN. Engineered biosynthesis of bacteriochlorophyll *b* in *Rhodobacter sphaeroides*. Biochimica et Biophysica Acta (BBA)-Bioenergetics. 2014 Oct 1;1837(10):1611–6.

28. Yanisch-Perron C, Vieira J, Messing J. Improved M13 phage cloning vectors and host strains: nucleotide sequences of the M13mpl8 and pUC19 vectors. Gene. 1985 Jan 1;33(1):103–19.

29. Thoma S, Schobert M. An improved Escherichia coli donor strain for diparental mating. FEMS microbiology letters. 2009 May 1;294(2):127–32.

30. Schäfer A, Tauch A, Jäger W, Kalinowski J, Thierbach G, Pühler A. Small mobilizable multi-purpose cloning vectors derived from the Escherichia coli plasmids pK18 and pK19: selection of defined deletions in the chromosome of Corynebacterium glutamicum. Gene. 1994 Jul 22;145(1):69–73.

31. Tikh IB, Held M, Schmidt-Dannert C. BioBrickTM compatible vector system for protein expression in Rhodobacter sphaeroides. Applied microbiology and biotechnology. 2014 Apr;98(7):3111–9.

32. Canniffe DP, Thweatt JL, Chew AG, Hunter CN, Bryant DA. A paralog of a bacteriochlorophyll biosynthesis enzyme catalyzes the formation of 1, 2-dihydrocarotenoids in green sulfur bacteria. Journal of Biological Chemistry. 2018 Sep 1;293(39):15233–42.

33. Magdaong NC, Niedzwiedzki DM, Goodson C, Blankenship RE. Carotenoid-to-bacteriochlorophyll energy transfer in the LH1–RC core complex of a bacteriochlorophyll b containing purple photosynthetic bacterium Blastochloris viridis. The Journal of Physical Chemistry B. 2016 Jun 16;120(23):5159–71.

34. Resnick SM, Madigan MT. Isolation and characterization of a mildly thermophilic nonsulfur purple bacterium containing bacteriochlorophyll b. FEMS microbiology letters. 1989 Nov 1;65(1-2):165–70.

35. Howarth NA, Rosenow J. Banning the bulb: Institutional evolution and the phased ban of incandescent lighting in Germany. Energy Policy. 2014 Apr 1;67:737–46.

36. Niedzwiedzki DM, Swainsbury DJ, Canniffe DP, Hunter CN, Hitchcock A. A photosynthetic antenna complex foregoes unity carotenoid-to-bacteriochlorophyll energy transfer efficiency to ensure photoprotection. Proceedings of the National Academy of Sciences. 2020 Mar 24;117(12):6502–8.

37. Wöhri AB, Katona G, Johansson LC, Fritz E, Malmerberg E, Andersson M, Vincent J, Eklund M, Cammarata M, Wulff M, Davidsson J. Light-induced structural changes in a photosynthetic reaction center caught by Laue diffraction. Science. 2010 Apr 30;328(5978):630–3.

38. Jones MR, Visschers RW, Van Grondelle R, Hunter CN. Construction and characterization of a mutant of Rhodobacter sphaeroides with the reaction center as the sole pigment-protein complex. Biochemistry. 1992 May 1;31(18):4458–65.

39. Canniffe DP, Hitchcock A. Photosynthesis. Carotenoids in photosynthesis: structure and biosynthesis. In: Jez J, ed. Encyclopedia of biological chemistry III, 3rd edn. 2021 (pp. 163–185). Elsevier, Oxford

40. Koyama Y, Fujii R. Cis-trans carotenoids in photosynthesis: configurations, excited-state properties and physiological functions. In The photochemistry of carotenoids 1999 (pp. 161–188). Springer, Dordrecht.

41. Malhotra HC, Britton G, Goodwin TW. Occurrence of 1, 2-dihydro-carotenoids in Rhodopseudomonas viridis. Journal of the Chemical Society D: Chemical Communications. 1970(2):127–8.

42. Imhoff JF, Rahn T, Künzel S, Neulinger SC. Phylogeny of anoxygenic photosynthesis based on sequences of photosynthetic reaction center proteins and a key enzyme in bacteriochlorophyll biosynthesis, the chlorophyllide reductase. Microorganisms. 2019 Nov 19;7(11):576.

43. Bylina EJ, Ohgi KA, Nute T, Cerniglia V, O’Neal S, Weaver P. A system for site-directed mutagenesis of the photosynthetic apparatus in Blastochloris viridis. Biotechnology et alia. 2002;9:1–23.

44. Thweatt JL, Canniffe DP, Bryant DA. Biosynthesis of chlorophylls and bacteriochlorophylls in green bacteria. Advances in botanical research. 2019 Jan 1;90:35–89.

45. Alberti M, Burke DH, Hearst JE. Structure and sequence of the photosynthesis gene cluster. In Anoxygenic photosynthetic bacteria 1995 (pp. 1083–1106). Springer, Dordrecht.

46. Madigan MT, Resnick SM, Kempher ML, Dohnalkova AC, Takaichi S, Wang-Otomo ZY, Toyoda A, Kurokawa K, Mori H, Tsukatani Y. Blastochloris tepida, sp. nov., a thermophilic species of the bacteriochlorophyll b-containing genus Blastochloris. Archives of Microbiology. 2019 Dec;201(10):1351–9.

47. Kyndt JA, Montano Salama D, Meyer TE. Genome sequence of the alphaproteobacterium Blastochloris sulfoviridis DSM 729, which requires reduced sulfur as a growth supplement and contains bacteriochlorophyll b. Microbiology Resource Announcements. 2020 Apr 30;9(18):e00313–20.

48. Stevens B. Water in the atmosphere. Phys. Today. 2013;66(6):29.

49. National Renewable Energy Laboratory (NREL), U.S. Department of Energy, Reference Air Mass 1.5 Spectra (2003);https://www.nrel.gov/grid/solar-resource/spectra-am1.5.html.

50. Francia F, Wang J, Venturoli G, Melandri BA, Barz WP, Oesterhelt D. The reaction center-LH1 antenna complex of Rhodobacter sphaeroides contains one PufX molecule which is involved in dimerization of this complex. Biochemistry. 1999 May 25;38(21):6834–45.

51. Komenda J, Sobotka R, Nixon PJ. Assembling and maintaining the photosystem II complex in chloroplasts and cyanobacteria. Current opinion in plant biology. 2012 Jun 1;15(3):245–51.

52. Niederman RA. Development and dynamics of the photosynthetic apparatus in purple phototrophic bacteria. Biochimica et Biophysica Acta (BBA)-Bioenergetics. 2016 Mar 1;1857(3):232–46.

53. Tank M, Thiel V, Ward DM, Bryant DA. A panoply of phototrophs: an overview of the thermophilic chlorophototrophs of the microbial mats of alkaline siliceous hot springs in Yellowstone National Park, WY, USA. Modern topics in the phototrophic prokaryotes. 2017:87–137.

54. Imhoff JF, Trüper HG. Ectothiorhodospira halochloris sp. nov., a new extremely halophilic phototrophic bacterium containing bacteriochlorophyll b. Archives of Microbiology. 1977 Aug;114(2):115–21.

55. Imhoff JF, Trüper HG. Ectothiorhodospira abdelmalekii sp. nov., a new halophilic and alkaliphilic phototrophic bacterium. Zentralblatt für Bakteriologie Mikrobiologie und Hygiene: I. Abt. Originale C: Allgemeine, angewandte und ökologische Mikrobiologie. 1981 Oct 1;2(3):228–34.

56. Steiner R, Scheer H. Characterisation of a B800/1020 antenna from the photosynthetic bacteria Ectothiorhodospira halochloris and Ectothiorhodospira abdelmalekii. Biochimica et Biophysica Acta (BBA)-Bioenergetics. 1985 May 31;807(3):278–84.

57. Wagner-Huber R, Brunisholz RA, Bissig I, Frank G, Suter F, Zuber H. The primary structure of the antenna polypeptides of Ectothiorhodospira halochloris and Ectothiorhodospira halophila: Four core-type antenna polypeptides in E. halochtoris and E. halophila. European Journal of biochemistry. 1992 May;205(3):917–25.

58. Tsukatani Y, Hirose Y, Harada J, Yonekawa C, Tamiaki H. Unusual features in the photosynthetic machinery of Halorhodospira halochloris DSM 1059 revealed by complete genome sequencing. Photosynthesis research. 2019 Jun;140(3):311–9.

59. Crouch LI, Jones MR. Cross-species investigation of the functions of the Rhodobacter PufX polypeptide and the composition of the RC–LH1 core complex. Biochimica et Biophysica Acta (BBA)-Bioenergetics. 2012 Feb 1;1817(2):336–52.

60. Kimura Y, Nojima S, Nakata K, Yamashita T, Wang XP, Takenaka S, Akimoto S, Kobayashi M, Madigan MT, Wang-Otomo ZY, Yu LJ. Electrostatic charge controls the lowest LH1 Qy transition energy in the triply extremophilic purple phototrophic bacterium, Halorhodospira halochloris. Biochimica et Biophysica Acta (BBA)-Bioenergetics. 2021 Nov 1;1862(11):148473.

61. Tani K, Kanno R, Makino Y, Hall M, Takenouchi M, Imanishi M, Yu LJ, Overmann J, Madigan MT, Kimura Y, Mizoguchi A. Cryo-EM structure of a Ca2+-bound photosynthetic LH1-RC complex containing multiple αβ-polypeptides. Nature communications. 2020 Oct 2;11(1):1–9.

