## Supplementary Material for "The role of the γ subunit in the photosystem of the lowest-energy phototrophs"

**SUPPLEMENTARY TABLES & FIGURES**

| Strain/Plasmid | Properties | Source |
|---|---|---|
| <u>*E. coli*</u> | | |
| JM109 | Cloning strain for pK18*mobsacB* and pBBRBB constructs | Promega |
| ST18 (DSM 22074) | Conjugative strain for pK18*mobsacB* and pBBRBB constructs | DSMZ |
| <u>*Blc. viridis*</u> | | |
| WT | DSM-133 | DSMZ |
| ΔLH1$\gamma_{1–3}$ | Replacement of BVIR_1786–1788 with *aadA* from pSRA81 in WT, *Sp*$^R$ | This study |
| ΔLH1$\gamma_{1–4}$ | Replacement of BVIR_2981 with *bla* from pET-3a in ΔLH1$\gamma_{1–3}$, *Sp*$^R$ *Amp*$^R$ | This study |
| ΔLH1$\gamma_{1–4}$ + LH1$\gamma_1$ | ΔLH1$\gamma_{1–4}$ harbouring pBBRBB-P*puf*$^{Bv}$[LH1$\gamma_1$], *Sp*$^R$ *Amp*$^R$ *Km*$^R$ | This study |
| ΔLH1$\gamma_{1–4}$ + LH1$\gamma_4$ | ΔLH1$\gamma_{1–4}$ harbouring pBBRBB-P*puf*$^{Bv}$[LH1$\gamma_4$], *Sp*$^R$ *Amp*$^R$ *Km*$^R$ | This study |
| <u>Plasmid</u> | | |
| pK18*mobsacB* | Allelic exchange vector, *Km*$^R$ | J. Armitage*, [2] |
| pBBRBB-P*puf*$_{843\text{-}1200}$-DsRed | Purple bacterial expression vector carrying the *Rhodobacter sphaeroides puf* promoter, *Km*$^R$ | Addgene, [3] |
| pBBRBB-P*puf*$^{Bv}$ | Replacement of promoter in pBBRBB-P*puf*$_{843\text{-}1200}$-DsRed with equivalent from *Blc. viridis*, *Km*$^R$ | This study |
| pBBRBB-P*puf*$^{Bv}$[LH1$\gamma_1$] | BVIR_1786 cloned downstream of promoter in pBBRBB-P*puf*$^{Bv}$ | This study |
| pBBRBB-P*puf*$^{Bv}$[LH1$\gamma_4$] | BVIR_2981 cloned downstream of promoter in pBBRBB-P*puf*$^{Bv}$ | This study |
| pSRA81 | Source of *aadA* cassette, *Sp*$^R$ | [4] |
| pET-3a | Source of *bla* cassette, *Amp*$^R$ | Novagen |

**Supplementary Table 1. List of strains and plasmids described in this study**
* Department of Biochemistry, University of Oxford, South Parks Road, Oxford OX1 3QU, U.K.


[1] Simon R, Priefer U, Pühler A (1983) A broad host range mobilization system for *in vivo* genetic engineering: transposon mutagenesis in Gram negative bacteria. *Nat Biotechnol* **1**:784–791

[2] Schäfer A, Tauch A, Jäger W, Kalinowski J, Thierbach G, Pühler A. (1994). Small mobilizable multi-purpose cloning vectors derived from the Escherichia coli plasmids pK18 and pK19: selection of defined deletions in the chromosome of Corynebacterium glutamicum. *Gene* 145:69–73

[3] Tikh IB, Held M, Schmidt-Dannert C. (2014) BioBrick™ compatible vector system for protein expression in *Rhodobacter sphaeroides*. *Appl Microbiol Biotechnol* **98**:3111–3119

[4] Canniffe DP, Thweatt JL, Chew AG, Hunter CN, Bryant DA. (2018) A paralog of a bacteriochlorophyll biosynthesis enzyme catalyzes the formation of 1, 2-dihydrocarotenoids in green sulfur bacteria. *J Biol Chem* **293**:15233–15242

| Primer | Sequence (5’-3’) | Cleavage site |
|---|---|---|
| LH1$\gamma_{1-3}$UpF | CCGGAATTCCCTTGAACCAGGCCTCCTCGCC | EcoRI |
| LH1$\gamma_{1-3}$UpR | CCAAAAAAACAGTCATAACAAGCCATCGTTGGTCCTCTCATGACGGGTC | |
| LH1$\gamma_{1-3}$DownF | CACCAAGGTAGTCGGCAAATAAGAATTGTCGGGTCCGGCCCCTATCG | |
| LH1$\gamma_{1-3}$DownR | CCCAAGCTTGGTGTTTGCCACCGCCATCGTCCTTG | HindIII |
| aadAF | ATGGCTTGTTATGACTGTTTTTTTGG | |
| aadAR | TTATTTGCCGACTACCTTGGTG | |
| LH1$\gamma_{1-3}$CheckF | CCAGGGCGTAATCCTCGGTGTC | |
| LH1$\gamma_{1-3}$CheckR | GCCAAGCTCCGCACCACGG | |
| LH1$\gamma_4$UpF | GAGTCTAGACCTTGGCTCCACCAAATTTCTTTTGCC | XbaI |
| LH1$\gamma_4$UpR | CGCACATTTCCCCGAAAAGTGCCGCTTTCTTCATTTTGCAGACTCC | |
| LH1$\gamma_4$DownF | CCTCACTGATTAAGCATTGGTAACTGCTAGTGACACGGTTTCCGGCC | |
| LH1$\gamma_4$DownR | CCCAAGCTTCTACGACCAGATCGCGGTCTCC | HindIII |
| blaF | GGCACTTTTCGGGGAAATGTGCG | |
| blaR | CAGTTACCAATGCTTAATCAGTGAGG | |
| LH1$\gamma_4$CheckF | GAATGGCATTCAAGAGGTCAGG | |
| LH1$\gamma_4$CheckR | GGCTTCCACTTCATGAACAAGG | |
| Ppuf$^{Bv}$F | GCTCTAGAGCTGATCCTCGACCATGATCG | XbaI |
| Ppuf$^{Bv}$R | CGAGATCTACCCTCATCAATGCGGGCC | BglII |
| LH1$\gamma_1$BBF | GCAGATCTATGAAACTTTCAGCTATTCTTG | BglII |
| LH1$\gamma_1$BBR | CTGACTAGTTCAACGATAGGTCAGCGCAATC | SpeI |
| LH1$\gamma_4$BBF | GCAGATCTATGAAGAAAGCATCTGCAATC | BglII |
| LH1$\gamma_4$BBR | CTGACTAGTCTAGTTGTAAACGAAGGCAATC | SpeI |

**Supplementary Table 2. List of primers used in this study**

Restriction enzyme cleavage sites used for cloning are underlined in the primer sequence.

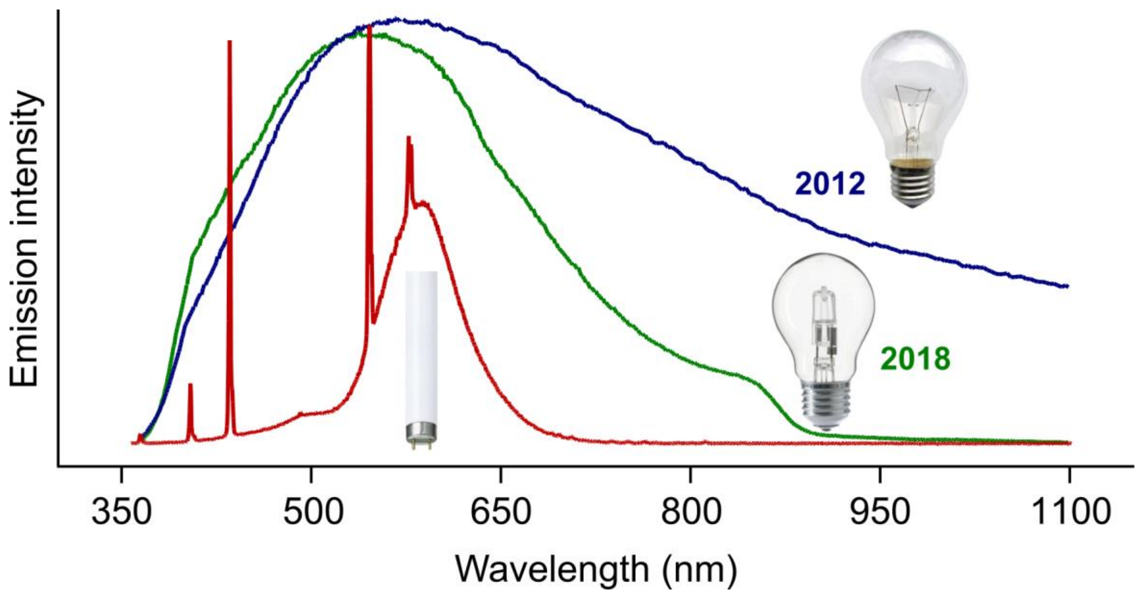


**Figure S1. Emission spectra of bulbs used for growth of *Blc. viridis* strains.** The year in which production of incandescent (blue) and halogen (green) bulbs were banned in the EU are labelled.

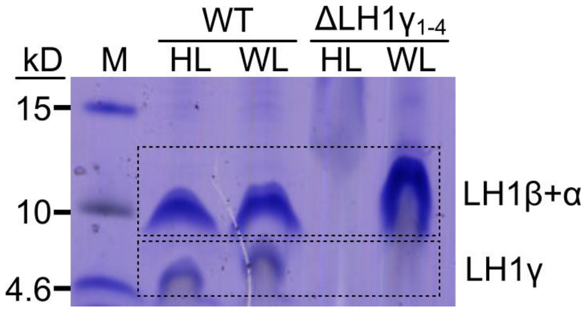


**Figure S2. Electrophoretic separation of LH1 components on a TRIS-tricine gel.**

A

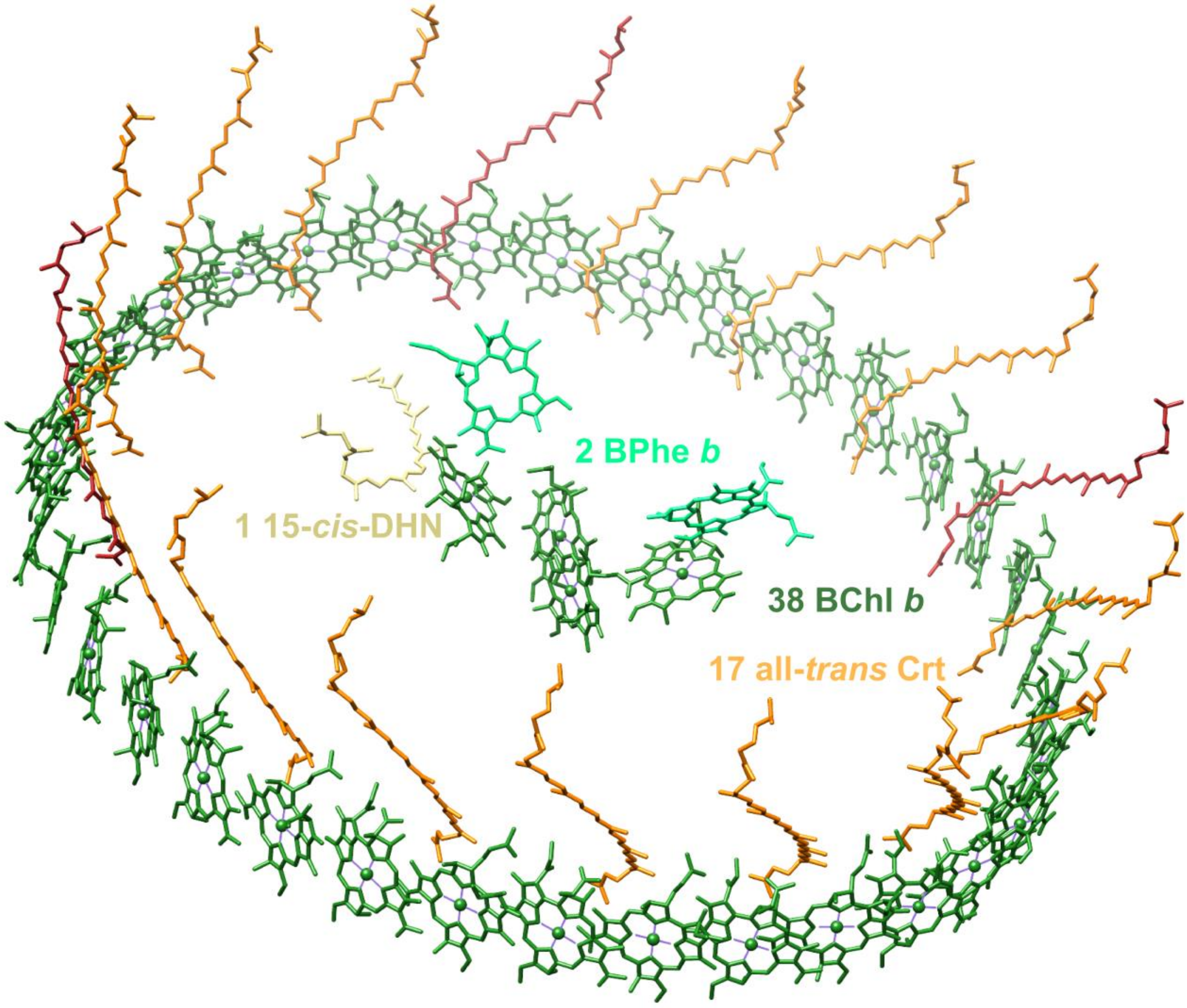


B

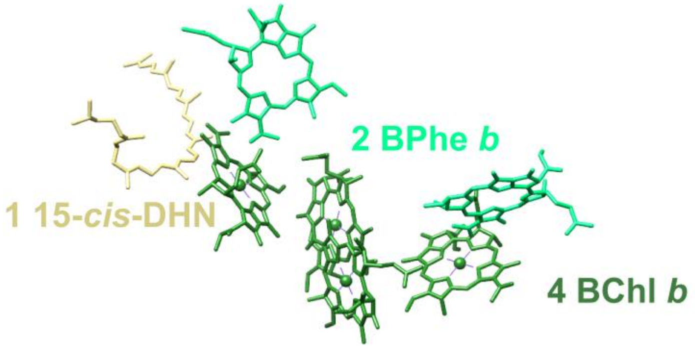


**Figure S3.** Arrangement and number of pigments in A) RC–LH1 and B) RC of *Blc. viridis*. Phytyl tails have been removed from BChls and BPhes for clarity. The 17 all-trans carotenoids in LH1 are coloured in orange (neurosporene species) or red (lycopene species) according to their approximate abundance in WT cells.
